# Genome-wide association studies reveal novel loci associated with carcass and body measures in goats (*Capra hircus*)

**DOI:** 10.1101/2025.03.24.644862

**Authors:** Linyun Zhang, Yixin Duan, Shengnan Zhao, Houmo Yu, Jipan Zhang, Naiyi Xu, Yongju Zhao

**Affiliations:** College of Animal Science and Technology, Southwest University, Chongqing 400715, China; Chongqing Key Laboratory of Herbivore Science, Chongqing 400715, China; Chongqing Key Laboratory of Forage & Herbivore, Chongqing 400715, China

**Keywords:** Goat, Whole-genome resequencing, GWAS, Growth trait, Carcass trait

## Abstract

Elucidating the genetic architecture of local goat populations is crucial for improving breeding strategies and conservation efforts. In southwestern China, there are numerous local goat resources, featuring rich phenotypic and genetic variations. Nevertheless, in-depth research on their production performance is scarce. Therefore, in this study, whole-genome 7× resequencing was performed on 776 individuals of six goat breeds (Dazu Black Goat, Yudong Black Goat, Banjiao Goat, Hechuan White Goat, Chuandong White Goat, and Youzhou Black Goat) in Chongqing to deeply investigate their population structure, genetic relationships, and heritability, and to identify significant genes related to growth traits and carcass traits. The mapping of significant genes for growth and carcass traits identified novel genetic loci, including a significant association at position 63,248,396 within *ASIP* on chromosome 13, which was linked to both traits, indicating a potential pleiotropic effect. Additional candidate genes for growth traits included *CXCL14, PTGFR*, and *DPYD* (body weight); *CCSER1, HMCN1*, and *MYOM1* (body length); *MALRD1* and *ASIP* (withers height); and *MEIS2* and *NLRC5* (chest circumference). For carcass traits, key candidate genes included *ASIP, ATP5G1, COX3* and *BCAR3*. Functional enrichment analysis highlighted the calcium signaling pathway, glycan biosynthesis, and cytokine receptor interactions as key biological pathways influencing these traits. These findings provide novel insights into the genetic mechanisms underlying economically important traits in Chongqing local goats. The identification of *ASIP* as a pleiotropic gene, along with other newly discovered loci, offers valuable targets for genetic improvement and selective breeding programs, enhancing the productivity and sustainability of local goat populations.

## Introduction

China is the world’s leading producer and consumer of sheep and goat products, with the highest numbers of sheep and goat inventory, slaughter, and meat production globally[1]. Chongqing, China, is home to six local goat breeds, including the Dazu Black Goat, Yudong Black Goat, Banjiao Goat, Hechuan White Goat, Chuandong White Goat, and Youzhou Black Goat (Figure 1). These breeds exhibit unique coat colors and adaptability, and they are widely distributed across different regions of Chongqing. These local goats are known for their excellent meat quality, tolerance to rough feed, and strong environmental adaptability, making them an important component of the local livestock industry.

**Figure 1.**
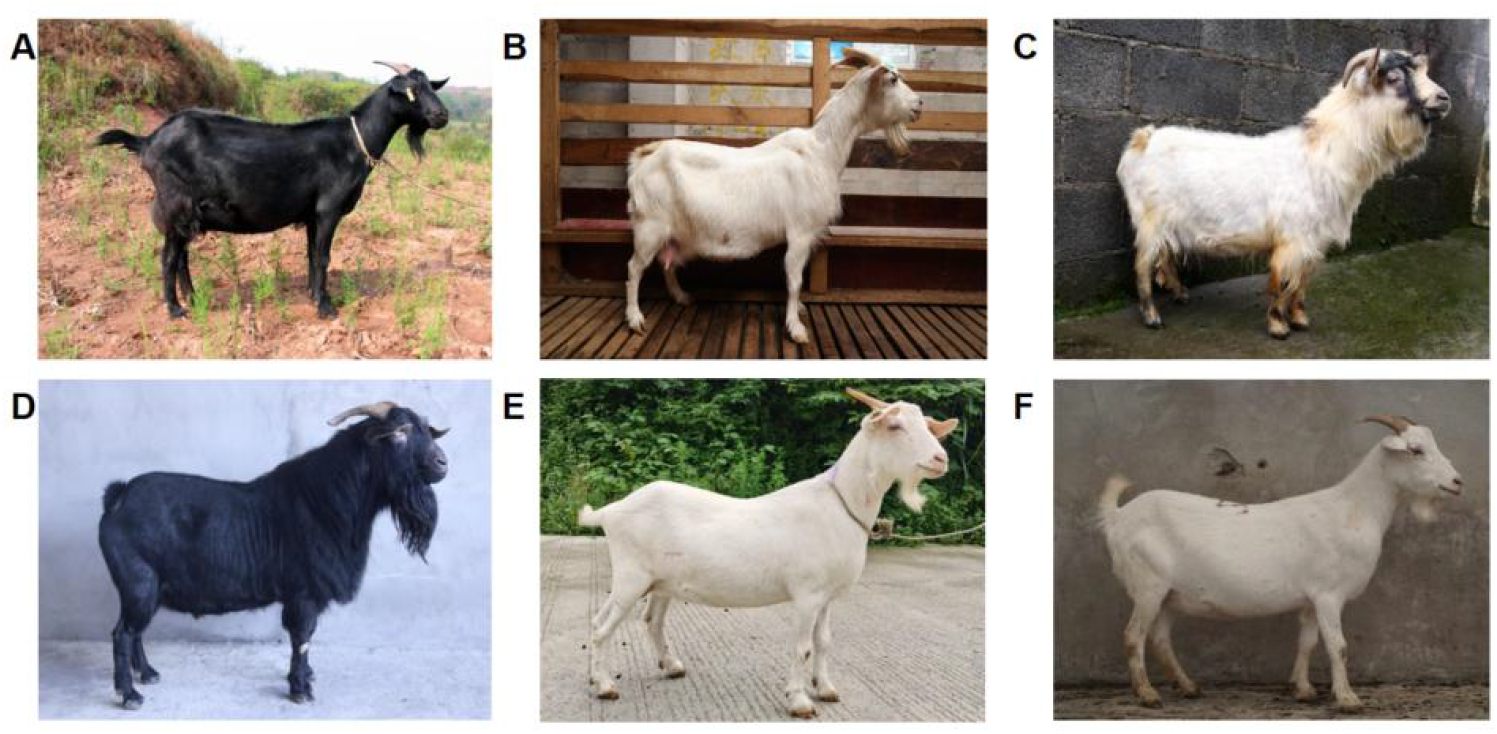
Pictures of six local goat breeds in Chongqing Note: A is Dazut black goat; B is Banjiao goat; C is Youzhou black goat; D is Yudong black goat; E is Hechuan white goat; F is Chuandong white goat.

Strengthening the technological self-reliance and control over breeding resources in the goat industry is crucial for the development of the meat goat sector and the broader livestock industry. With the increasing availability of high-depth whole-genome resequencing data and advancements in sequencing technologies, the identification of candidate genes associated with economically important traits has become more precise [2]. Comprehensive analysis of genome-wide single nucleotide polymorphisms (SNPs) allows for the accurate mapping of key genomic regions linked to target traits, providing a solid scientific foundation for genetic improvement and the advancement of goat breeding worldwide [3].

Growth traits and carcass traits are important economic traits in meat goat, reflecting their growth and development and being directly related to production. GWAS has been widely applied to study various economic traits, and genes such as *BTA14, LYN, LYPLA1, MRPL15, BTA6, and BTA16* have been found to be potentially associated with growth and carcass traits [4,5]. Further understanding of the genetic mechanisms underlying these traits can aid in genetic improvement, enhancing breeding efficiency and economic benefits.

This study combines population structure analysis and genetic correlation analysis with GWAS technology to analyze the growth and carcass traits of Chongqing local goats. The aim is to identify candidate genes significantly associated with these traits and explore their potential biological functions and mechanisms through gene function annotation and pathway analysis.

## Materials and methods

### Experimental animals

The breed name, quantity and sampling location of the Chongqing local goats used in this study are listed in Table 1.

**Table 1.**
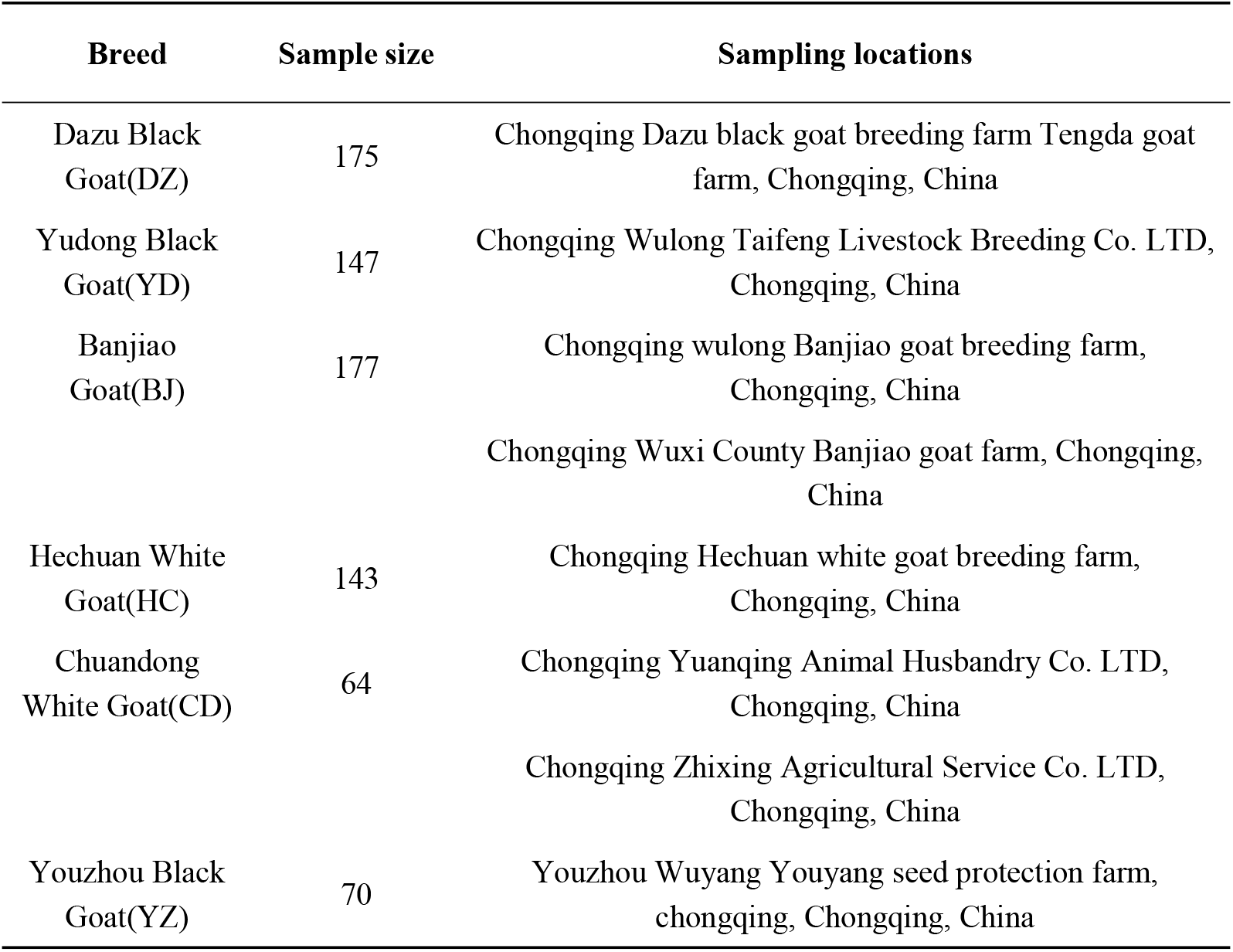
Sampling information of six Chongqing local goat breeds.

### Whole-genome resequencing and quality control

Using the DNBSEQ-T7 platform (Complete Genomics and MGI Tech, Shenzhen, China), 776 samples were re-sequenced for 7 × by Using the DNBSEQ-T7 platform (Complete Genomics and MGI Tech, Shenzhen, China). The raw sequencing data were filtered to remove adapter sequences or low-quality reads by Plink1.9. Then, the clean reads were then aligned to the goat reference genome (ARS1.2) using the BWA software. Subsequently, the BAM files were sorted using Picard and Samtools. SNPs detection was performed using GATK, and the necessary sample information was extracted from the VCF files. Finally, the sequencing data underwent quality control using PLINK, with the following criteria:

1. Removal of SNPs with a missing rate greater than 10%.
2. Removal of loci with a minor allele frequency (MAF) less than 1%.
3. Removal of loci with a Hardy-Weinberg equilibrium p-value less than 1×10^−6^.

### Genetic relationship and population structure analysis of Chongqing local goats

Based on whole-genome SNP data, population structure for ancestral components with K=2 to K=6 was calculated using Admixture [6], and visualized with the package “pophelper” of R [7,8]. The NJ phylogenetic tree was constructed using a genetic distance matrix generated by Plink1.9 [9], with the tree file created by FastTree (http://www.microbesonline.org/fasttree/) and visualized using FigTree v1.4.4 (http://tree.bio.ed.ac.uk/software/figtree/). The A kinship heatmap was generated in R using the “pheatmap” package [8] after constructing the genetic relationship matrix (G) with Gemma [10]. LD decay analysis and visualization were performed using PopLDdecay [11]. The PCA analysis based on quality-controlled SNP data was conducted using Plink, and the results were visualized with the “CMplot” package [12] in R.

### SNP heritability estimation

The Genome-wide Restricted Maximum Likelihood (GREML) method is utilized for estimating SNP heritability. The mixed linear model (MLM), initially applied by Yang [13] for SNP heritability estimation, assumes that all SNPs have non-zero effects and that the effect sizes (*β*) follow a normal distribution:

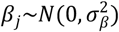

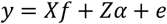

In this equation, *y* represents the phenotypic trait, *f* denotes fixed effects (non-genetic), *a* signifies random additive effects, and *e* stands for residual effects. *X* and *Z* are the corresponding design matrices.

The SNP heritability 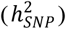 estimate is given by: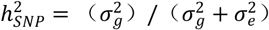, where the additive genetic variance 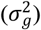 is defined as: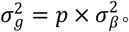

### Genome-wide association study

In this study, GWAS between SNPs and growth traits as well as carcass traits by conducted using a mixed linear model in the GEMMA software. The model is specified as follows:

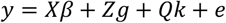

In this equation, *y* represents the phenotypic data, *β* is the vector of fixed effects excluding SNP and kinship (such as PCA, sex, etc.), *g* is the vector of SNP effects, *k* represents the vector of random effects (kinship), and *e* is the vector of residual effects. *X, Z*, and *Q* are the corresponding design matrices.

The GWAS results were visualized using Manhattan plot and QQ plot generated with the qqman [14] and “CMplot” packages in R.

### Gene Function annotation and pathway analysis

The Gene Ontology (GO) database classifies gene functions into three main categories: Cellular Component (CC), Molecular Function (MF), and Biological Process (BP). KEGG is a database for metabolic pathways. Candidate genes were identified based on their positions in the ARS1.2, using the upstream and downstream 200kb distances of significant loci as the interval. SNP loci were annotated to corresponding genes using bedtools software. GO and KEGG functional enrichment analyses were performed using the DAVID online tool (https://david.ncifcrf.gov/), and the results were visualized using the “ggplot2” package [15] in R.

## Result

### Genetic relationship and population structure analysis of Chongqing local goats Population structure analysis of Chongqing local goats

We performed population structure analysis of six Chongqing local breeds (DZ, YD, BJ, CD, HC, YZ) with ancestors numbers from K=2 to K=6 (Figure 2A). As the value of K increases, each population shows an increasingly complex genetic composition, when K ranges from 4 to 6 the populations already display more diverse genetic components and significant internal genetic diversity. Based on the complexity of genetic components and the stability of population structure, when K=4 and K=5, the genetic components of different populations show a clear separation. While they clearly reveal the population structure, they also maintain reasonable model complexity, making the results more biologically explanatory, K=4 and K=5 are likely the most appropriate model choices. These values can adequately explain the genetic backgrounds and historical migration patterns of the populations.

**Figure 2.**
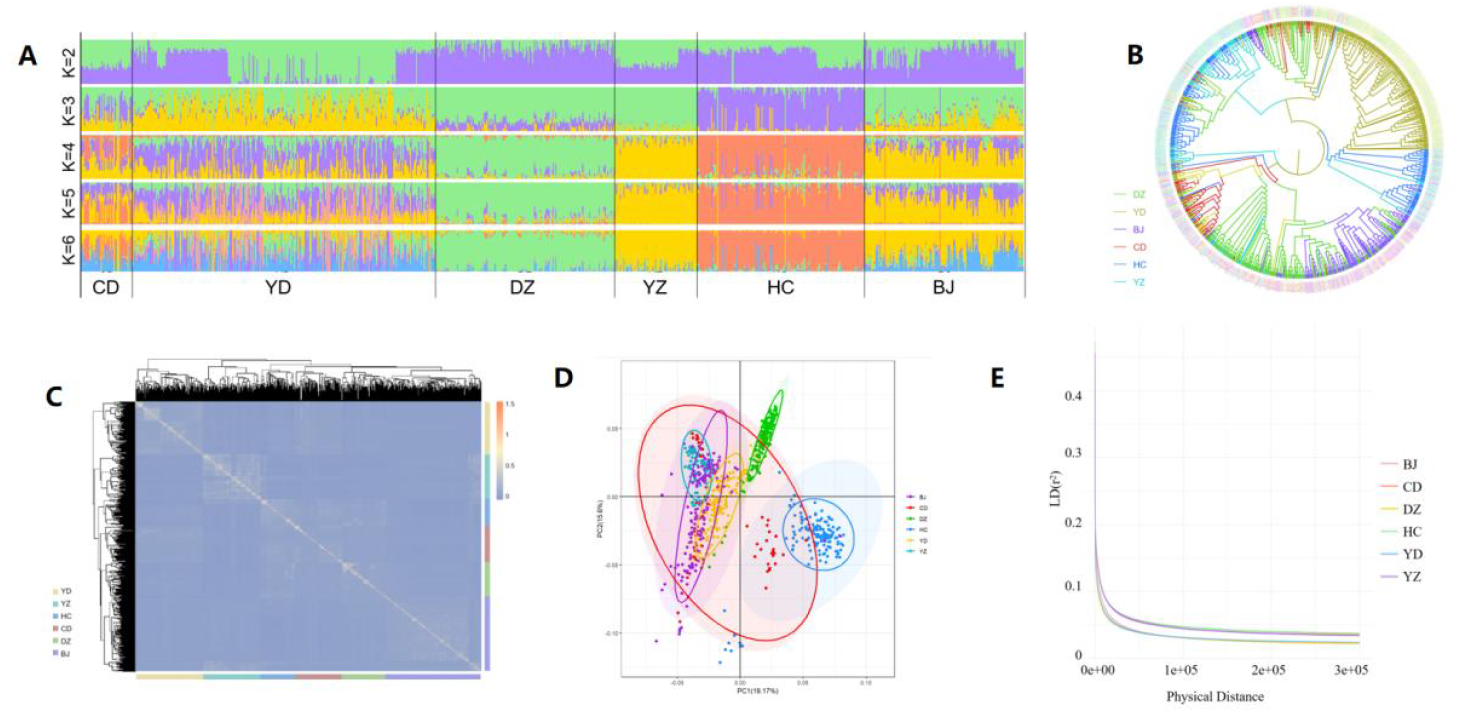
Analysis of population structure in Chongqing local goat breeds Note: A is the population structure analysis result by admixture; B is the Neighbor-Joining (NJ) phylogenetic tree; C is the Genomic relationship matrix; D is the Principal Component Analysis of overall population; E is the LD decay; DZ represents the Dazu Black Goat; YD represents the Yudong Black Goat; BJ represents the Banjiao Goat; CD represents the Chuandong White Goat; HC represents the Hechuan White Goat; YZ represents the Youzhou Black Goat.

The genetic distance matrix was computed, and a Neighbor-Joining (NJ) phylogenetic tree was subsequently constructed with FastTree. The phylogenetic tree of Chongqing local goat breeds was visualized by FigTree (Figure 2B). The tree showing that each breed of Chongqing local goat exhibits a unique and distinct clade.

Using whole-genome SNP data from 175 Dazu Black Goats, 147 Yudong Black Goats, 177 Banjiao Goats, 143 Hechuan White Goats, 64 Chuandong White Goats, and 70 Youzhou Black Goats to construct the genetic matrix (G matrix). Based on the matrix G, the genomic relationship heatmap (Figure 2C) was then generated by using the “pheatmap” package in R. The color scale in the Figure 2C represents different breeds of Chongqing local goat, and the genomic relationship coefficients between most individuals range from 0 to 0.5. It is evident that there is high similarity within each breed, while the similarity between different breeds is relatively low.

Principal component analysis (PCA) was performed using the genomic matrix (G) of Chongqing local goats (Figure 2D). Distinct clusters were formed along the PC1 (19.17%) and PC2 (15.6%) axes. The HC formed distinct, independent clusters, whereas the remaining breeds exhibited partial overlap with each other. This observation suggests substantial genetic differentiation among the populations, alongside some shared genetic similarities.

The linkage disequilibrium (LD) decay plots for the different populations show that the LD decay trends are generally similar across the populations (Figure 2E). The mean correlation coefficients r^2^ between loci for DZ, BJ, YD, CD, HC, and YZ are 0.032, 0.033, 0.033, 0.047, 0.049, and 0.047, respectively; when r^2^ decays to 0.1, the physical distances are approximately 3.39 kb, 5.48 kb, 3.97 kb, 8.91 kb, 8.48 kb, and 8.49 kb, respectively. The trend of maintaining linkage disequilibrium between closely spaced genetic markers is similar among the populations, indicating similar genetic structures and evolutionary histories.

### Correlation analysis of growth traits and estimation of genetic parameters in Chongqing local goats

Based on the SNP information of the whole genome, the heritability and genetic correlation estimation of goat growth traits were studied, and combined with phenotypic correlation, the growth trait correlation analysis table of Chongqing local goats was drawn as shown below(Table 2). It can be seen from the table that the growth traits of goats have low to moderate heritability, and there are significant positive correlations among the growth traits.

**Table 2.**
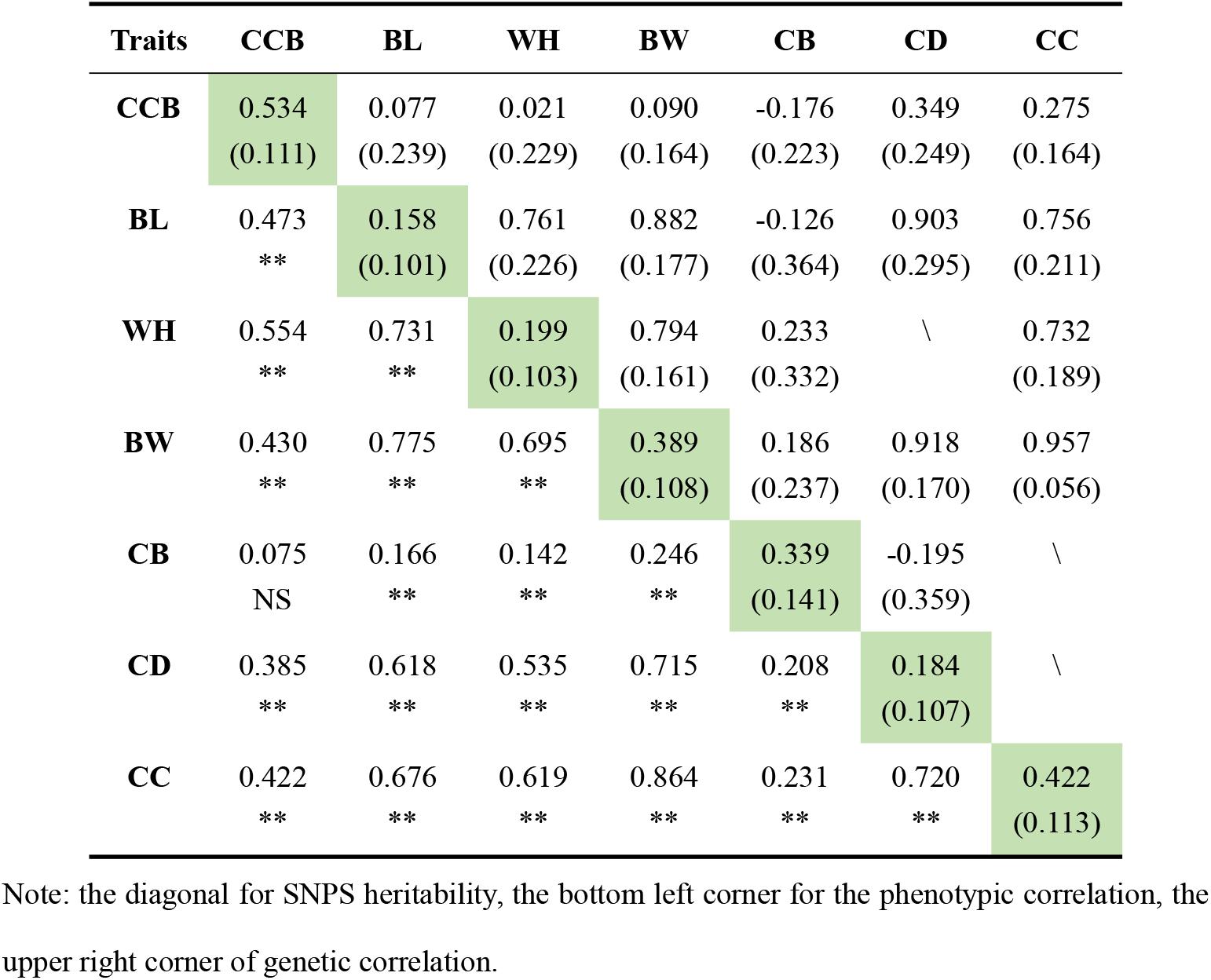
The growth character correlation analysis of Chongqing local goats

| Traits | CCB | BL | WH | BW | CB | CD | CC |
| --- | --- | --- | --- | --- | --- | --- | --- |
| **CCB** | 0.534<br>(0.111) | 0.077<br>(0.239) | 0.021<br>(0.229) | 0.090<br>(0.164) | -0.176<br>(0.223) | 0.349<br>(0.249) | 0.275<br>(0.164) |
| **BL** | 0.473<br>** | 0.158<br>(0.101) | 0.761<br>(0.226) | 0.882<br>(0.177) | -0.126<br>(0.364) | 0.903<br>(0.295) | 0.756<br>(0.211) |
| **WH** | 0.554<br>** | 0.731<br>** | 0.199<br>(0.103) | 0.794<br>(0.161) | 0.233<br>(0.332) | \ | 0.732<br>(0.189) |
| **BW** | 0.430<br>** | 0.775<br>** | 0.695<br>** | 0.389<br>(0.108) | 0.186<br>(0.237) | 0.918<br>(0.170) | 0.957<br>(0.056) |
| **CB** | 0.075<br>NS | 0.166<br>** | 0.142<br>** | 0.246<br>** | 0.339<br>(0.141) | -0.195<br>(0.359) | \ |
| **CD** | 0.385<br>** | 0.618<br>** | 0.535<br>** | 0.715<br>** | 0.208<br>** | 0.184<br>(0.107) | \ |
| **CC** | 0.422<br>** | 0.676<br>** | 0.619<br>** | 0.864<br>** | 0.231<br>** | 0.720<br>** | 0.422<br>(0.113) |

Note: the diagonal for SNPS heritability, the bottom left corner for the phenotypic correlation, the upper right corner of genetic correlation.

### GWAS of growth traits in Chongqing local goats

The GWAS was performed on growth traits (BW, WH, BL, CC, CD) across six goat breeds, the results were visualized using R (Figure 3A-E), with a significance threshold set at p < 1.52×10^-07^. Based on site 200kb interval of upstream and downstream of the gene annotation.

**Figure 3.**
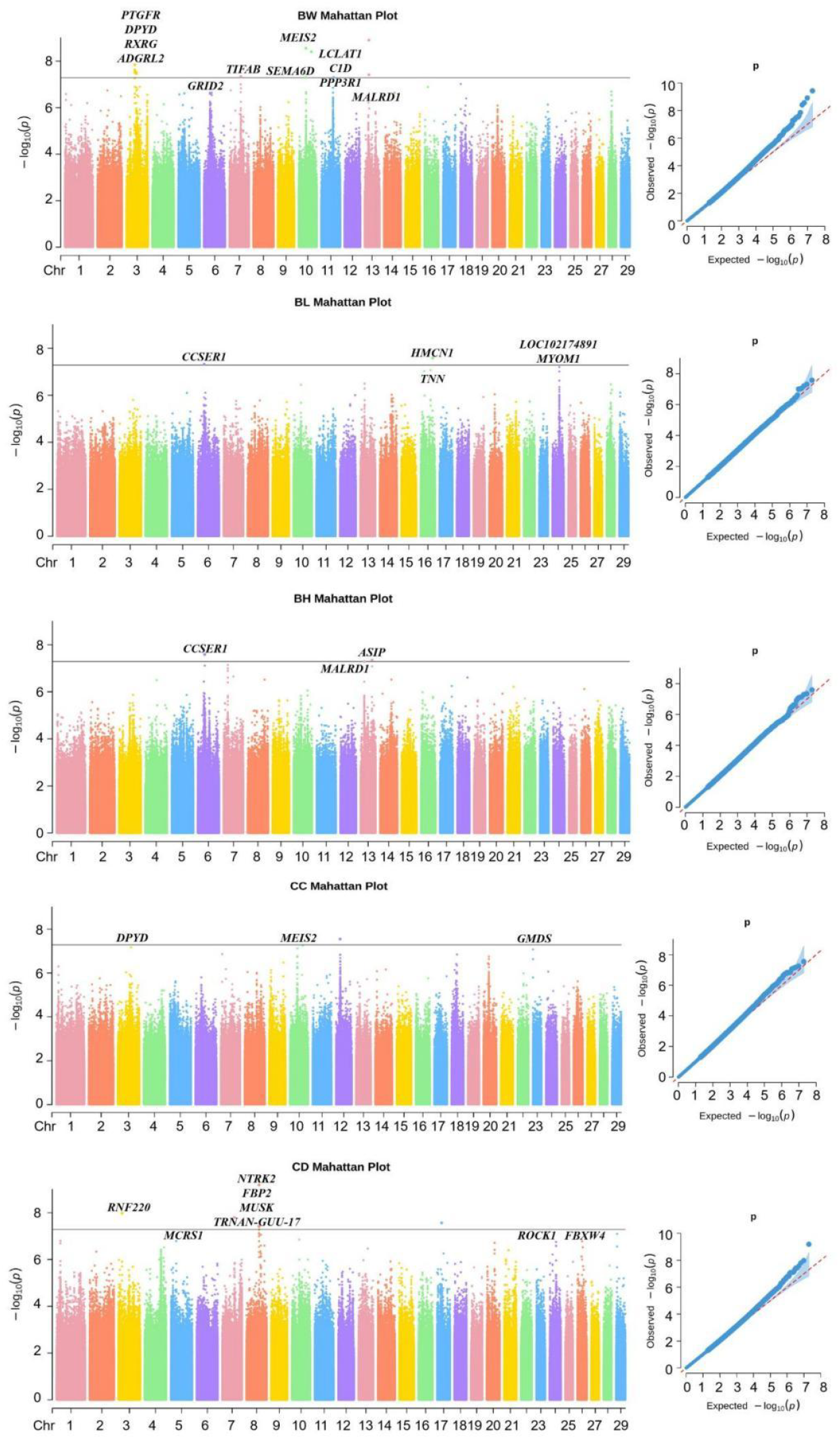
Mahattan plots and QQ plots of GWAS results for growth traits of Chongqing local goats Note: A is body weight; B is body length; C is withers height; D is chest circumference; E is chest depth (*p* =1.52×10^-07^).

The GWAS for BW (Figure 3A) identified several potential candidate genes significantly associated with BW on chromosome 3, 7, 10, 11 and 13, including *CXCL14, PTGFR, DPYD* etc.. The GWAS for BL (Figure 3B) revealed significant associations on chromosome 6, 16, and 24, with candidate genes such as *CCSER1, HMCN1*, and *MYOM1*. For WH (Figure 3C), significant associations were found on chromosomes 6 and 13, with potential candidate genes including *CCSER1, MALRD1*, and *ASIP*. For CC (Figure 3D) indicated significant associations on chromosome 4, 10 and 23, with candidate genes such as *DPYD, MEIS2*, and *NLRC5*. Finally, the GWAS for CD (Figure 3E) identified significant associations on chromosomes 3 and 8, with potential candidate genes including *RNF220, MUSK*, and *EBP2*. The chromosome, location, p-value and nearby gene information of the significant SNPs for each growth traits are shown in Table S1.

### The GWAS for traits of carcass in three Chongqing local goat breed

In six local goat breeds from Chongqing, significant differences in growth traits such as body size and weight were observed among Dazu Black Goat, Yudong Black Goat, and Banjiao Goat. And 30 individuals of these three breeds from the population used for the GWAS on live growth traits were selected for carcass traits analysis.

#### PCA of three Chongqing local goat

For the three breeds, based on the PCA results (Figure 4A), add PC1, PC2 enough for assessing group structure, PC1, PC2 in subsequent correlation analysis as a covariate to join to the model.

**Figure 4.**
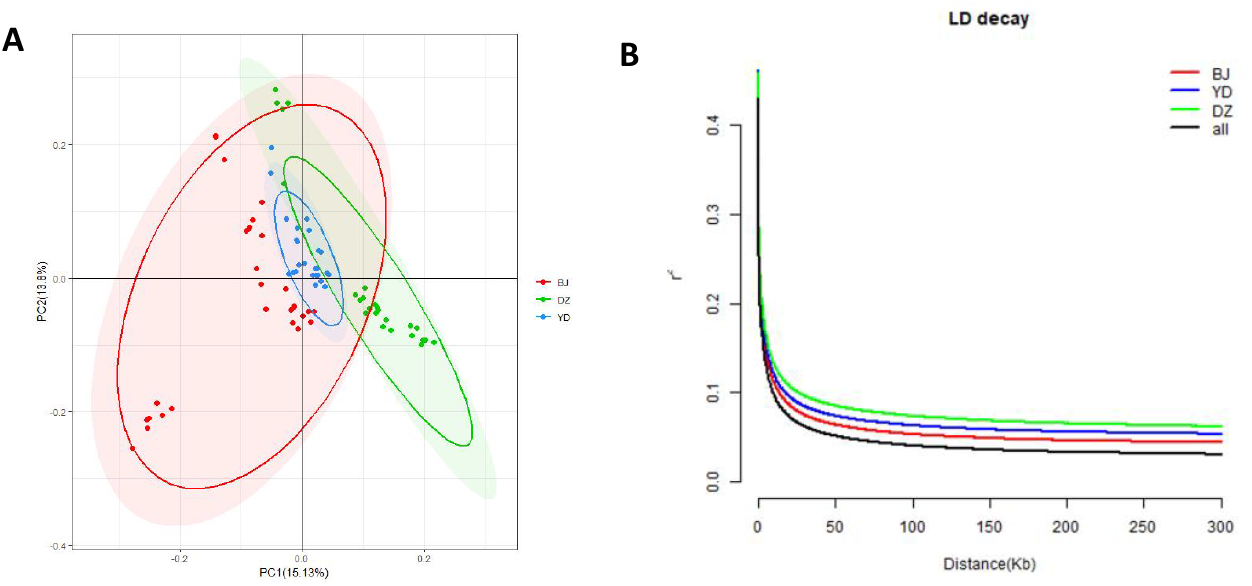
Visualization of principal component analysis and LD decayof multi-breed goats Note: A: PCA two-dimensional diagram red represents Banjiao goats, blue represents Yudong black goats, and green represents Dazu black goats; B: LD decay of mixed population, Banjiao Goat, Yudong Black Goat and Dazu Black Goat.

The LD decay plot (Figure 4B) showed that when r^2^ was 0.1, the decay distance of the mixed population was 9.5kb, and the decay distance of Banjiao goat, Yudong black goat and Dazu black goat was 13kb, 17.5kb and 26kb, respectively.

### Determination of slaughtered carcass traits

The experimental animals were fasted for 24h and water deprived for 2h before slaughter. The live weight (LW) before slaughter was weighed, and then lumbar muscle area (LMA), GR value (GR), carcass weight (CW), net meat weight (NMW), carcass percentage (CP), meat percentage (MP), and net carcass percentage (NCP) were measured. The determination method was based on “Technical Specification for the Determination of Production Performance of cotton and Goat” (NY/T1236-2006) as the reference standard. The formula is as follows:

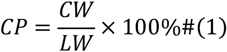

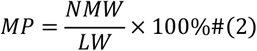

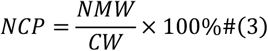

In three breeds and mixed population for LMA, GR, CW, NMW, CP, MP and NCP such as screening to the SNP loci and candidate genes (significant sites upstream and downstream LD decay distance interval) from Table S2, and Figure 5.

**Figure 5.**
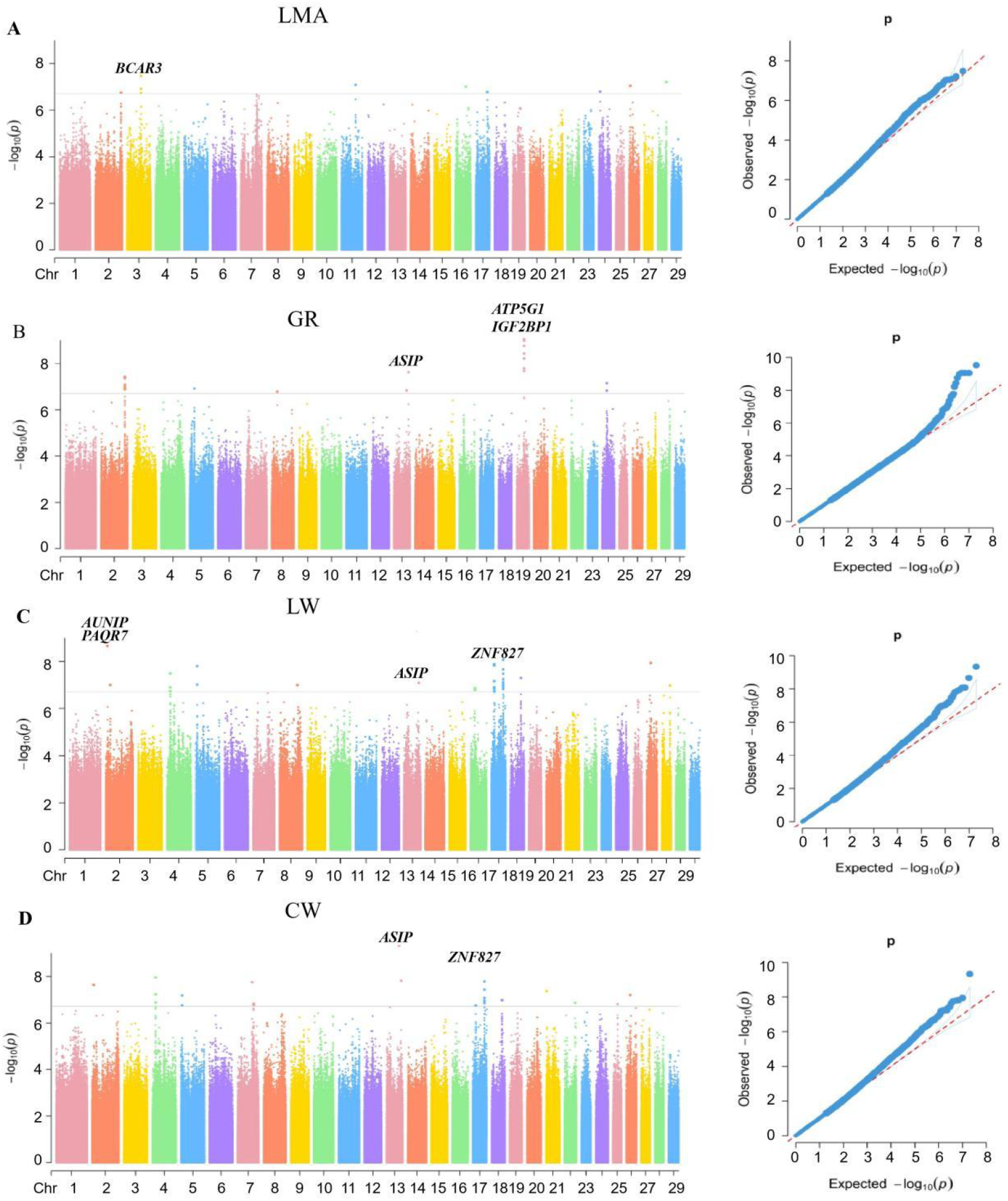
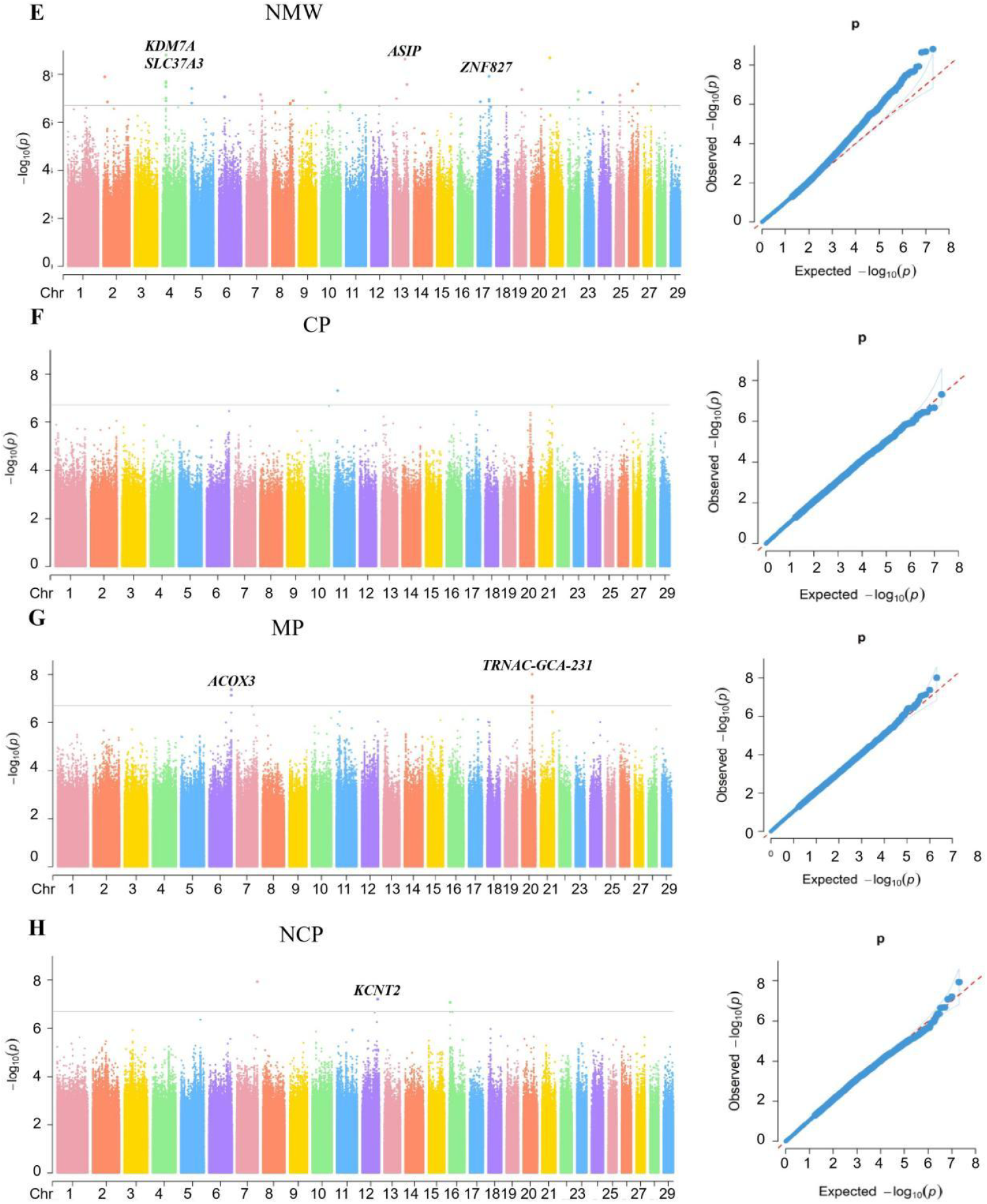
Manhattan plots and QQ plots of carcass traits of multi-breed goats Note: A: Lumbar muscle area; B: GR value; C: live weight before slaughter; D: carcass weight; E: net meat weight; F: Dressing percentage; G: meat percentage; H: net meat percentage (*p* = 1.96×10^-07^).

Based on site 200kb interval of upstream and downstream of the gene annotation. Through mixed population analysis, numerous genes including *ASIP, ATP5G1, ACOX3, BCAR3, TRNAC-GCA-231*, and *ZNF827* were identified as potential candidate genes associated with carcass traits in goats.

### Gene function annotation and pathway analysis

In growth traits of Chongqing local goats, the KEGG pathway enrichment analysis was performed on the significant genes associated (Figure 6A). The size of each bubble represents the number of genes involved in each pathway, with larger bubbles indicating a higher gene count. The color gradient of the bubbles reflects the p-value of enrichment, where blue signifies lower p-values, indicative of greater statistical significance, and red denotes higher p-values. Notable pathways identified include “Calcium signaling pathway,” which demonstrated significant gene involvement and relatively low p-values, suggesting a strong association with the observed growth traits. In addition, the Gene Ontology (GO) enrichment analysis was conducted on differentially expressed genes linked to growth traits in Chongqing local goats (Figure 6B). The bar plot is color-coded according to GO categories: GO_BP (Biological Process) in orange, GO_CC (Cellular Component) in blue, and GO_MF (Molecular Function) in green. The analysis revealed enrichment in 12 GO_CC entries, 4 GO_MF entries, and 14 GO_BP entries. Significant terms highlighted include “glutamatergic synapse” under the GO_CC category and “calcium ion binding” under the GO_MF category, emphasizing their crucial roles in regulating the growth traits of Chongqing goats.

**Fig6.**
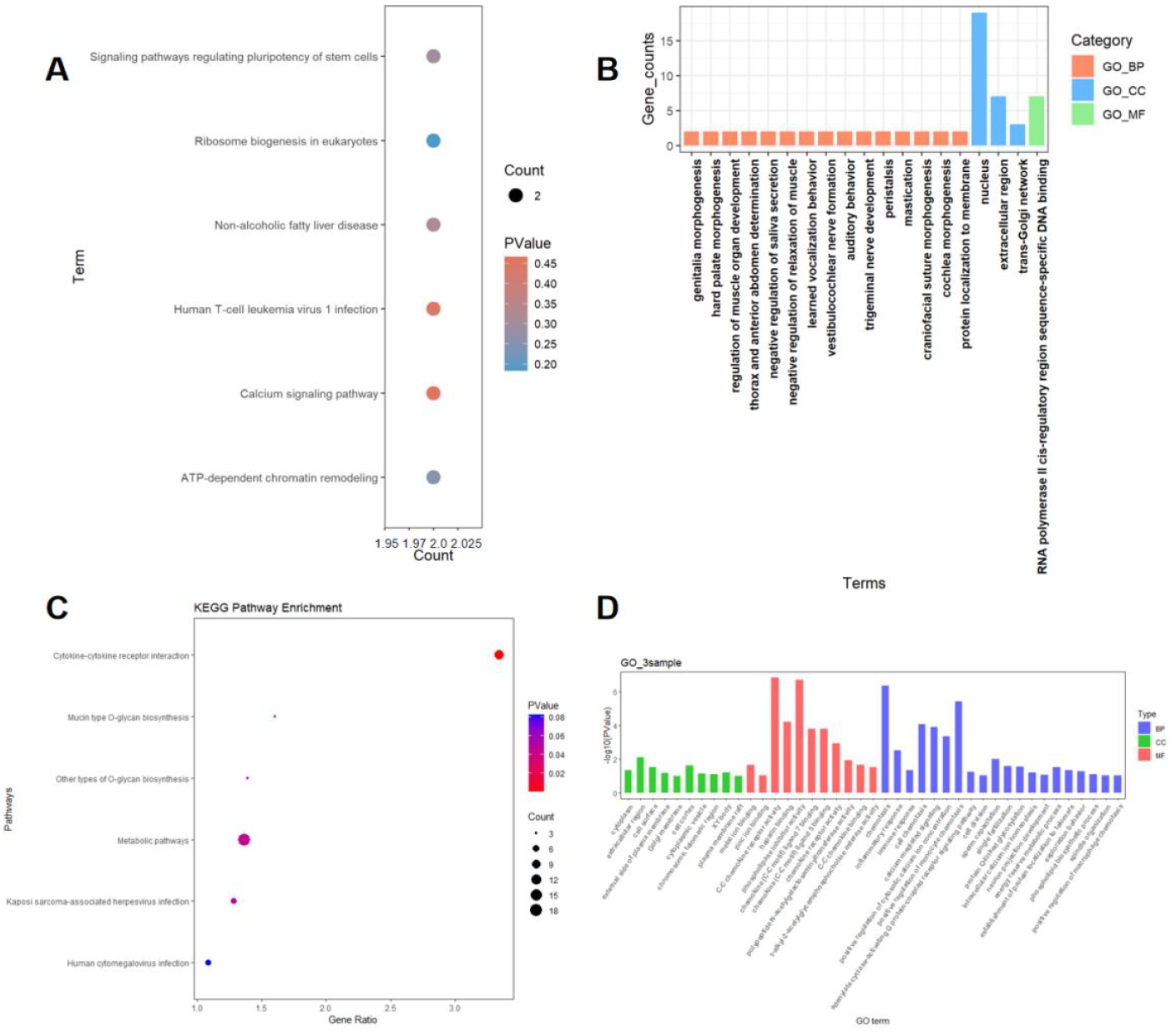
KEGG and Go results for significant genes of the coat color in Chongqing local goats Note: A represents the KEGG results for the candidate gene of growth traits; B represents the GO results for growth traits; C represents the KEGG results for carcass traits; D represents the GO results for carcass traits.

The GO and KEGG enrichment analysis of the candidate genes for carcass traits of goats in the three populations showed that 10 CC items, 11 MF items and 20 BP items were enriched, which were mainly related to cell components such as membranes, molecular functions such as chemokine bioactivity, energy metabolism and phospholipid biosynthesis and other biological processes, as shown in Fig 6D. Six pathways were co-enriched, mainly related to glycan biosynthesis, cytokine receptor interactions, and metabolic pathways, as shown in Fig 6C.

## Discussion

The comprehensive analysis of Chongqing local goats, encompassing six distinct breeds, revealed intricate genetic structures and relationships. Population structure analysis indicated significant intra-breed similarity and inter-breed differentiation, underscoring the unique genetic identities of each breed. Through GWAS, several candidate genes associated with key growth and carcass traits were identified, such as *ASIP, PTGFR, DPYD, MEIS2*, and *CCSER1*. These genes are involved in various biological processes, including growth regulation, metabolic pathways, skeletal development, and muscle formation, making them promising targets for genetic improvement.

The locus located at position 63,248,396 within the *ASIP* gene was identified in both growth and carcass traits. As an endogenous antagonist of the melanocortin receptor, ASIP is associated with melanin production. Previous studies have also found that *ASIP* is related to lipid metabolism. Xie et al. [16] identified ASIP as a candidate gene influencing lipid metabolism in bovine adipocytes. Using CRISPR/Cas9 technology to knock out the *ASIP* gene in bovine mammary epithelial cells, they found that *ASIP* knockout downregulated the expression of PPARγ, FASN, and SCD, affecting the saturation of fatty acids in milk. This indicates that ASIP plays a crucial role in regulating lipid metabolism in bMECs (bovine mammary epithelial cells). Additionally, Liu et al. [17] demonstrated through polymorphism and haplotype analysis of the ASIP gene that it is significantly associated with fat-related traits in Chinese Simmental cattle and has a significant correlation with carcass traits such as body weight and backfat thickness. The identification of the ASIP gene in relation to black coat color and its potential impact on growth traits suggests a pleiotropic effect, warranting further investigation to explore its dual role. Additionally, candidate genes such as *CCSER1* and *MYOM1*, associated with BL and BH, *MEIS2* associated with BW and CC, respectively, underscore the genetic basis underlying physical conformation traits in local goats. In addition, *CCSER1* and *GRID2* genes, which were significantly related to the growth traits of Chongqing local goats in this study, were located to be related to the growth traits of Red Angus in beef cattle and also had a certain correlation with the carcass traits of beef cattle [18,19]. The *CCSER1* gene has also been found to be associated with growth in sheep [20]. The *DPYD* gene was also found to be associated with goat growth in Guizhou black goats [21].

Functional enrichment analysis provided insights into the biological pathways involved, with the Calcium signaling pathway as significant for growth and carcass traits. Moreover, pathways related to glycan biosynthesis and cytokine receptor interactions indicate the involvement of immune and metabolic processes in carcass quality traits. These findings enhance our understanding of the molecular mechanisms governing growth and carcass traits, paving the way for the development of more targeted breeding programs.

Integrating these genetic insights into selective breeding strategies can help accelerate genetic progress and improve the productivity and economic value of Chongqing local goat breeds. By focusing on key genes and pathways, future breeding programs can be optimized to enhance growth performance, meat quality, and overall adaptation to local environmental conditions, contributing to the sustainability and profitability of goat farming in the region. Further research will be essential to validate these candidate genes and pathways and to uncover their precise roles in the genetic architecture of growth and carcass traits in goats.

## Conclusion

This study provides novel insights into the genetic architecture of Chongqing local goats by integrating population structure analysis, GWAS, and functional enrichment analyses. The large-scale, high-depth genomic dataset generated in this study serves as a valuable resource for future global research on goat genetics. Our findings highlight the genetic differentiation among breeds and identify key loci influencing growth and carcass traits. The identification of *ASIP* as a potential pleiotropic gene, along with other candidate genes such as *CCSER1* and *MEIS2*, suggests a complex genetic basis underlying economically important traits. Enrichment analysis further emphasizes the role of pathways involved in skeletal growth, fat metabolism, and immune regulation.

This dataset, derived from indigenous goat populations in Southwest China, provides unique insights into local adaptations and genetic diversity, offering a reference for comparative studies and precision breeding worldwide. The utilization of these genetic markers in future breeding programs has the potential to enhance meat production efficiency, adaptability, and genetic diversity in local goat populations. Further functional validation of the identified candidate genes and pathways is essential for translating genomic insights into practical applications in precision breeding.

## List of Abbreviations

**Abbreviation The full name of abbreviations**

GWAS: Genome-wide Association Studies
SNPs: Single Nucleotide Polymorphisms
DZ: Dazu Black Goat
YD: Yudong Black Goat
BJ: Banjiao Goat
HC: Hechuan White Goat
CD: Chuandong White Goat
YZ: Youzhou Black Goat
BW: Body Weight
BL: Body Length
WH: Withers Height
CC: Chest Circumference
CD: Chest Depth
CCB: Circumference of Cannon Bone
CB: Chest Breadth
LW: Live Weight
LMA: Lumbar Muscle Area
CW: Carcass Weight
NMW: Net Meat Weight
CP: Carcass Percentage
MP: Meat Percentage
NCP: Net Carcass Percentage
MAF: Minor Allele Frequency
GREML: Restricted Maximum Likelihood MLM Mixed Linear Model
PCA: Principal Component Analysis
LD: Linkage Disequilibrium

## Declarations

## Ethics approval and consent to participate

All animal management and experimental procedures followed the animal care protocols approved by the Ethics Committee of Southwest University (No. IACUC-20230417-02).

## Acknowledgments

This work was funded by the National Key Research and Development Program of China (No.2022YFD1300202), the Collection, Utilization and Innovation of Animal Resources by Research Institutes and Enterprises of Chongqing(No.Cqnyncw-kqlhtxm), Chongqing Modern Agricultural Industry Technology System (CQMAITS202313), and the National Training Program of Innovation and Entrepreneurship for Undergraduates (No. S202310635040). And we thank the Hefei advanced computing center for providing numerical computations resources for this paper.

## Consent for publication

Not applicable.

## CRediT authorship contribution statement

Linyun Zhang: Conceptualization, Methodology, Software, Investigation, Formal Analysis, Visualization, Writing - Original Draft; Yixin Duan: Software, Investigation, Formal Analysis, Writing - Original Draft; Shengnan Zhao: Visualization, Investigation; Houmo Yu: Investigation; Jipan Zhang: Writing - Review & Editing; Naiyi Xu:Conceptualization, Supervision, Writing - Review & Editing; Yongju Zhao: Conceptualization, Funding Acquisition, Resources, Supervision, Writing - Review & Editing.

## Attachments

**Table S1.**
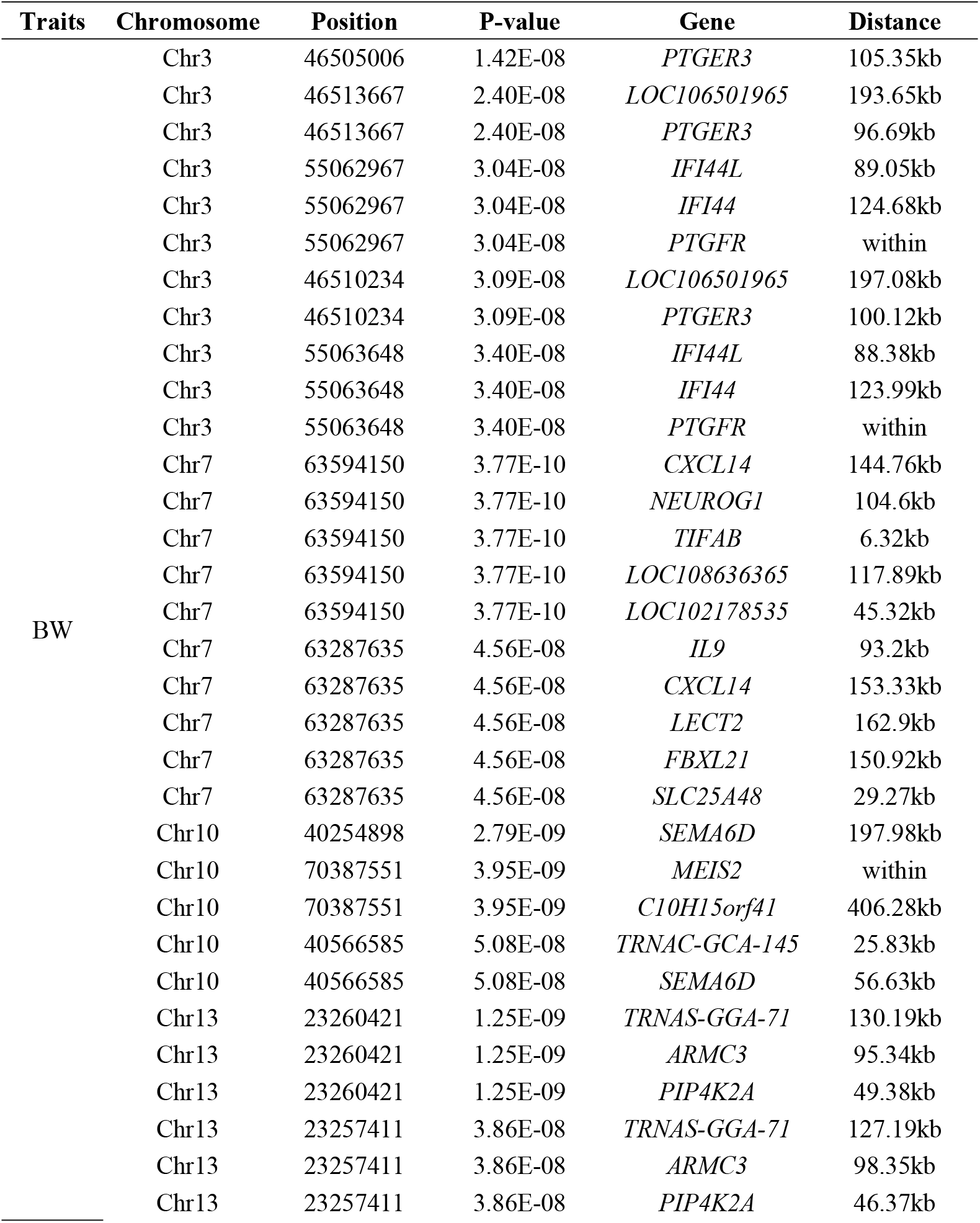
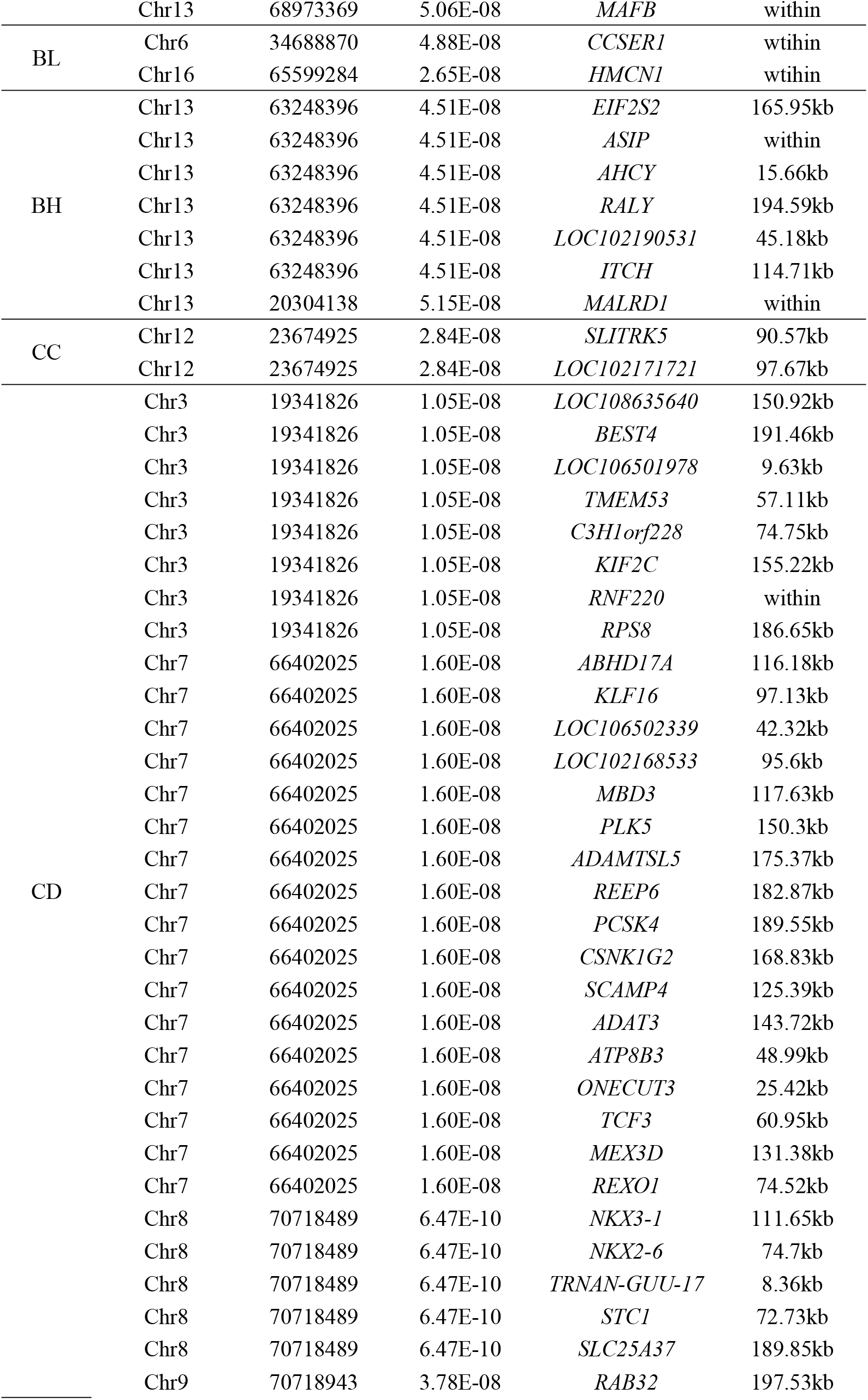
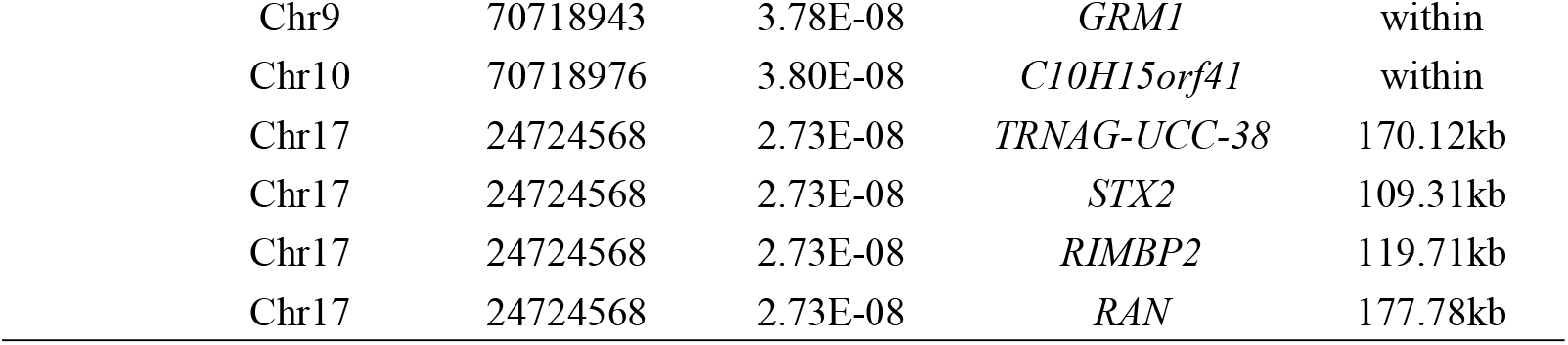
The genes about significant SNPs for growth traits in six breeds of Chongqing local goat

| Traits | Chromosome | Position | P-value | Gene | Distance |
| --- | --- | --- | --- | --- | --- |
| BW | Chr3 | 46505006 | 1.42E-08 | *PTGER3* | 105.35kb |
|  | Chr3 | 46513667 | 2.40E-08 | *LOC106501965* | 193.65kb |
|  | Chr3 | 46513667 | 2.40E-08 | *PTGER3* | 96.69kb |
|  | Chr3 | 55062967 | 3.04E-08 | *IFI44L* | 89.05kb |
|  | Chr3 | 55062967 | 3.04E-08 | *IFI44* | 124.68kb |
|  | Chr3 | 55062967 | 3.04E-08 | *PTGFR* | within |
|  | Chr3 | 46510234 | 3.09E-08 | *LOC106501965* | 197.08kb |
|  | Chr3 | 46510234 | 3.09E-08 | *PTGER3* | 100.12kb |
|  | Chr3 | 55063648 | 3.40E-08 | *IFI44L* | 88.38kb |
|  | Chr3 | 55063648 | 3.40E-08 | *IFI44* | 123.99kb |
|  | Chr3 | 55063648 | 3.40E-08 | *PTGFR* | within |
|  | Chr7 | 63594150 | 3.77E-10 | *CXCL14* | 144.76kb |
|  | Chr7 | 63594150 | 3.77E-10 | *NEUROG1* | 104.6kb |
|  | Chr7 | 63594150 | 3.77E-10 | *TIFAB* | 6.32kb |
|  | Chr7 | 63594150 | 3.77E-10 | *LOC108636365* | 117.89kb |
|  | Chr7 | 63594150 | 3.77E-10 | *LOC102178535* | 45.32kb |
|  | Chr7 | 63287635 | 4.56E-08 | *IL9* | 93.2kb |
|  | Chr7 | 63287635 | 4.56E-08 | *CXCL14* | 153.33kb |
|  | Chr7 | 63287635 | 4.56E-08 | *LECT2* | 162.9kb |
|  | Chr7 | 63287635 | 4.56E-08 | *FBXL21* | 150.92kb |
|  | Chr7 | 63287635 | 4.56E-08 | *SLC25A48* | 29.27kb |
|  | Chr10 | 40254898 | 2.79E-09 | *SEMA6D* | 197.98kb |
|  | Chr10 | 70387551 | 3.95E-09 | *MEIS2* | within |
|  | Chr10 | 70387551 | 3.95E-09 | *C10H15orf41* | 406.28kb |
|  | Chr10 | 40566585 | 5.08E-08 | *TRNAC-GCA-145* | 25.83kb |
|  | Chr10 | 40566585 | 5.08E-08 | *SEMA6D* | 56.63kb |
|  | Chr13 | 23260421 | 1.25E-09 | *TRNAS-GGA-71* | 130.19kb |
|  | Chr13 | 23260421 | 1.25E-09 | *ARMC3* | 95.34kb |
|  | Chr13 | 23260421 | 1.25E-09 | *PIP4K2A* | 49.38kb |
|  | Chr13 | 23257411 | 3.86E-08 | *TRNAS-GGA-71* | 127.19kb |
|  | Chr13 | 23257411 | 3.86E-08 | *ARMC3* | 98.35kb |
|  | Chr13 | 23257411 | 3.86E-08 | *PIP4K2A* | 46.37kb |
|  | Chr13 | 68973369 | 5.06E-08 | *MAFB* | within |
| BL | Chr6 | 34688870 | 4.88E-08 | *CCSER1* | wtihin |
|  | Chr16 | 65599284 | 2.65E-08 | *HMCN1* | wtihin |
| BH | Chr13 | 63248396 | 4.51E-08 | *EIF2S2* | 165.95kb |
|  | Chr13 | 63248396 | 4.51E-08 | *ASIP* | within |
|  | Chr13 | 63248396 | 4.51E-08 | *AHCY* | 15.66kb |
|  | Chr13 | 63248396 | 4.51E-08 | *RALY* | 194.59kb |
|  | Chr13 | 63248396 | 4.51E-08 | *LOC102190531* | 45.18kb |
|  | Chr13 | 63248396 | 4.51E-08 | *ITCH* | 114.71kb |
|  | Chr13 | 20304138 | 5.15E-08 | *MALRD1* | within |
| CC | Chr12 | 23674925 | 2.84E-08 | *SLITRK5* | 90.57kb |
|  | Chr12 | 23674925 | 2.84E-08 | *LOC102171721* | 97.67kb |
| CD | Chr3 | 19341826 | 1.05E-08 | *LOC108635640* | 150.92kb |
|  | Chr3 | 19341826 | 1.05E-08 | *BEST4* | 191.46kb |
|  | Chr3 | 19341826 | 1.05E-08 | *LOC106501978* | 9.63kb |
|  | Chr3 | 19341826 | 1.05E-08 | *TMEM53* | 57.11kb |
|  | Chr3 | 19341826 | 1.05E-08 | *C3H1orf228* | 74.75kb |
|  | Chr3 | 19341826 | 1.05E-08 | *KIF2C* | 155.22kb |
|  | Chr3 | 19341826 | 1.05E-08 | *RNF220* | within |
|  | Chr3 | 19341826 | 1.05E-08 | *RPS8* | 186.65kb |
|  | Chr7 | 66402025 | 1.60E-08 | *ABHD17A* | 116.18kb |
|  | Chr7 | 66402025 | 1.60E-08 | *KLF16* | 97.13kb |
|  | Chr7 | 66402025 | 1.60E-08 | *LOC106502339* | 42.32kb |
|  | Chr7 | 66402025 | 1.60E-08 | *LOC102168533* | 95.6kb |
|  | Chr7 | 66402025 | 1.60E-08 | *MBD3* | 117.63kb |
|  | Chr7 | 66402025 | 1.60E-08 | *PLK5* | 150.3kb |
|  | Chr7 | 66402025 | 1.60E-08 | *ADAMTSL5* | 175.37kb |
|  | Chr7 | 66402025 | 1.60E-08 | *REEP6* | 182.87kb |
|  | Chr7 | 66402025 | 1.60E-08 | *PCSK4* | 189.55kb |
|  | Chr7 | 66402025 | 1.60E-08 | *CSNK1G2* | 168.83kb |
|  | Chr7 | 66402025 | 1.60E-08 | *SCAMP4* | 125.39kb |
|  | Chr7 | 66402025 | 1.60E-08 | *ADAT3* | 143.72kb |
|  | Chr7 | 66402025 | 1.60E-08 | *ATP8B3* | 48.99kb |
|  | Chr7 | 66402025 | 1.60E-08 | *ONECUT3* | 25.42kb |
|  | Chr7 | 66402025 | 1.60E-08 | *TCF3* | 60.95kb |
|  | Chr7 | 66402025 | 1.60E-08 | *MEX3D* | 131.38kb |
|  | Chr7 | 66402025 | 1.60E-08 | *REXO1* | 74.52kb |
|  | Chr8 | 70718489 | 6.47E-10 | *NKX3-1* | 111.65kb |
|  | Chr8 | 70718489 | 6.47E-10 | *NKX2-6* | 74.7kb |
|  | Chr8 | 70718489 | 6.47E-10 | *TRNAN-GUU-17* | 8.36kb |
|  | Chr8 | 70718489 | 6.47E-10 | *STC1* | 72.73kb |
|  | Chr8 | 70718489 | 6.47E-10 | *SLC25A37* | 189.85kb |
|  | Chr9 | 70718943 | 3.78E-08 | *RAB32* | 197.53kb |

|  |  |  |  |  |
| --- | --- | --- | --- | --- |
| Chr9 | 70718943 | 3.78E-08 | *GRM1* | within |
| Chr10 | 70718976 | 3.80E-08 | *C10H15orf41* | within |
| Chr17 | 24724568 | 2.73E-08 | *TRNAG-UCC-38* | 170.12kb |
| Chr17 | 24724568 | 2.73E-08 | *STX2* | 109.31kb |
| Chr17 | 24724568 | 2.73E-08 | *RIMBP2* | 119.71kb |
| Chr17 | 24724568 | 2.73E-08 | *RAN* | 177.78kb |

**Table S2.**
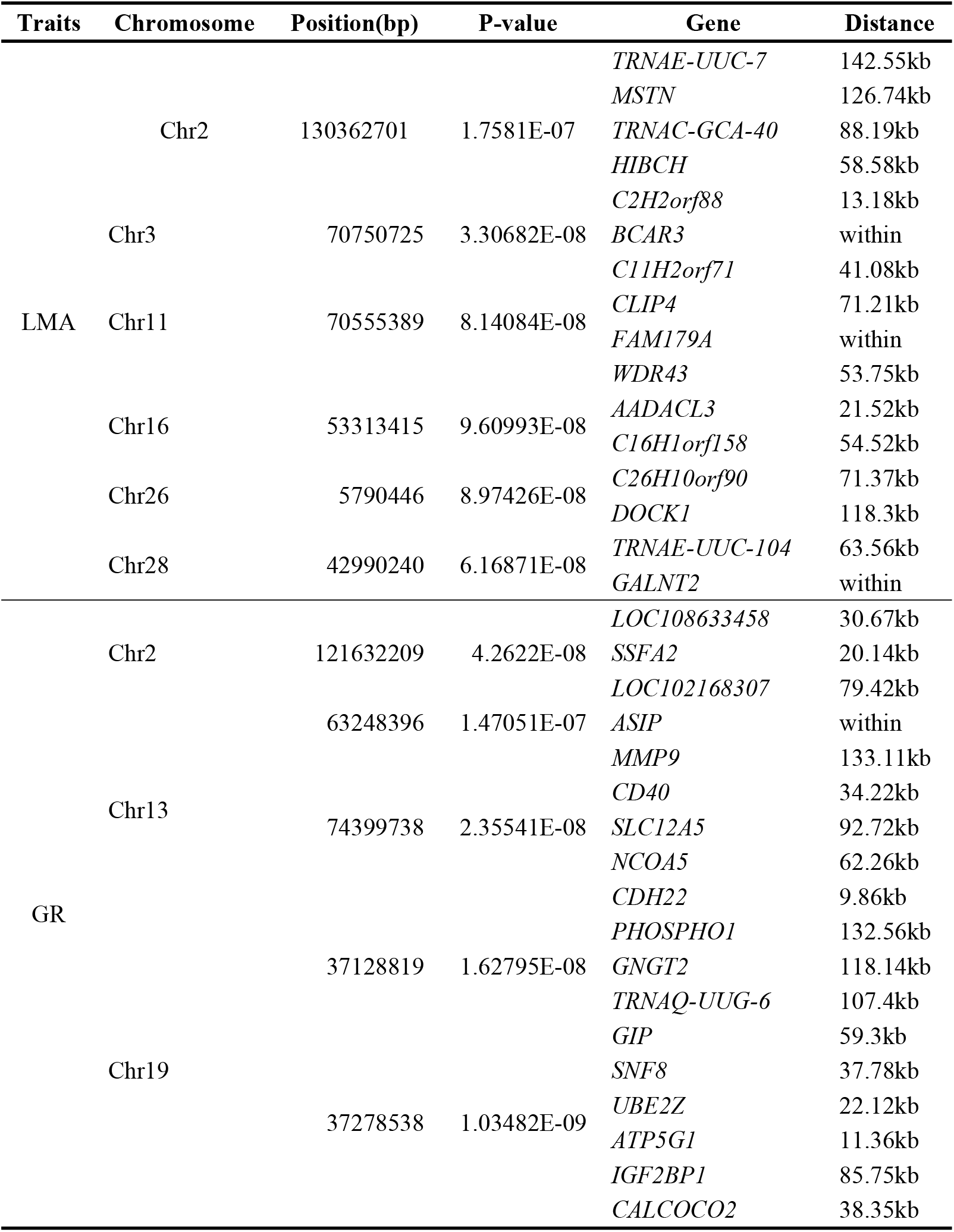
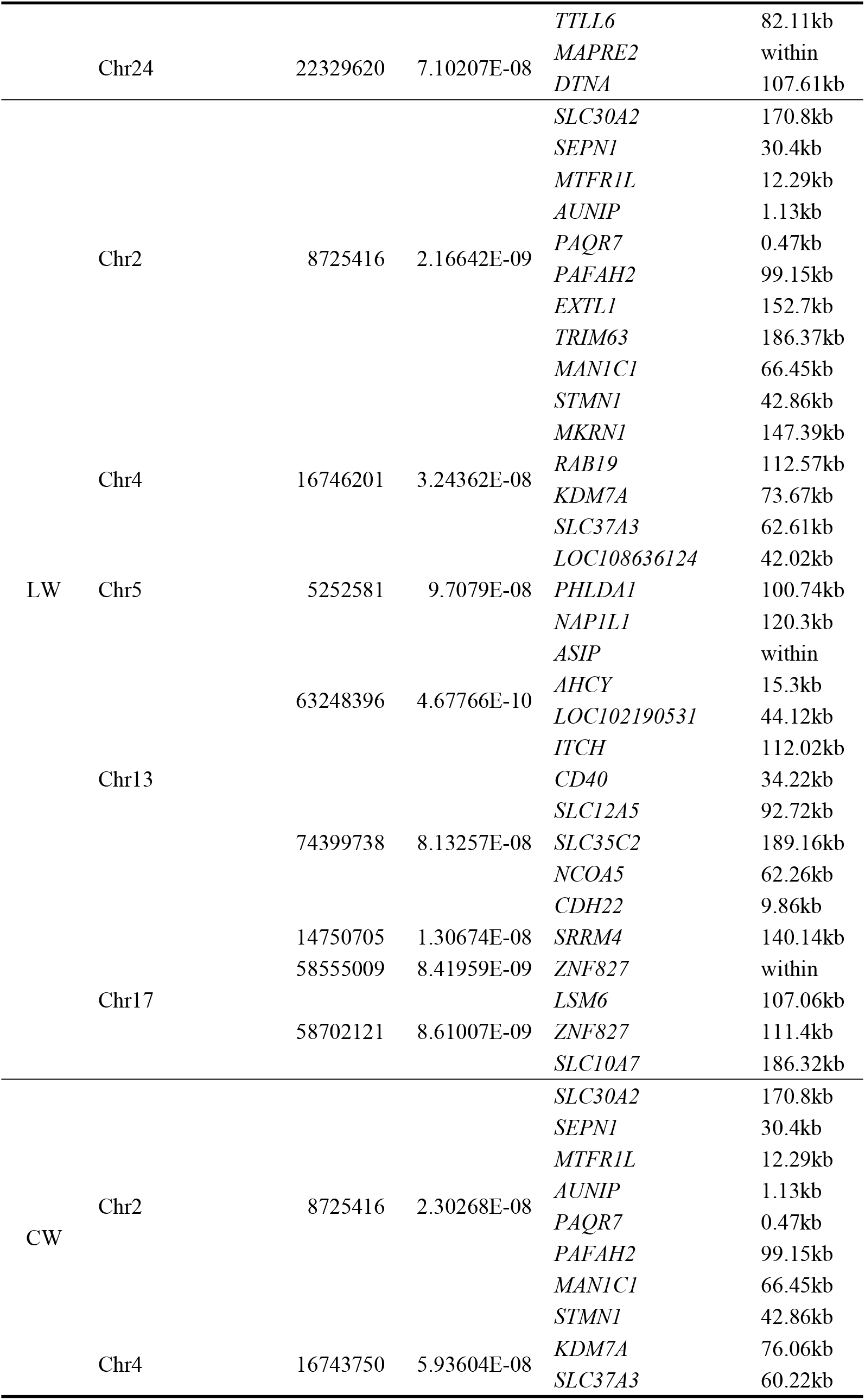
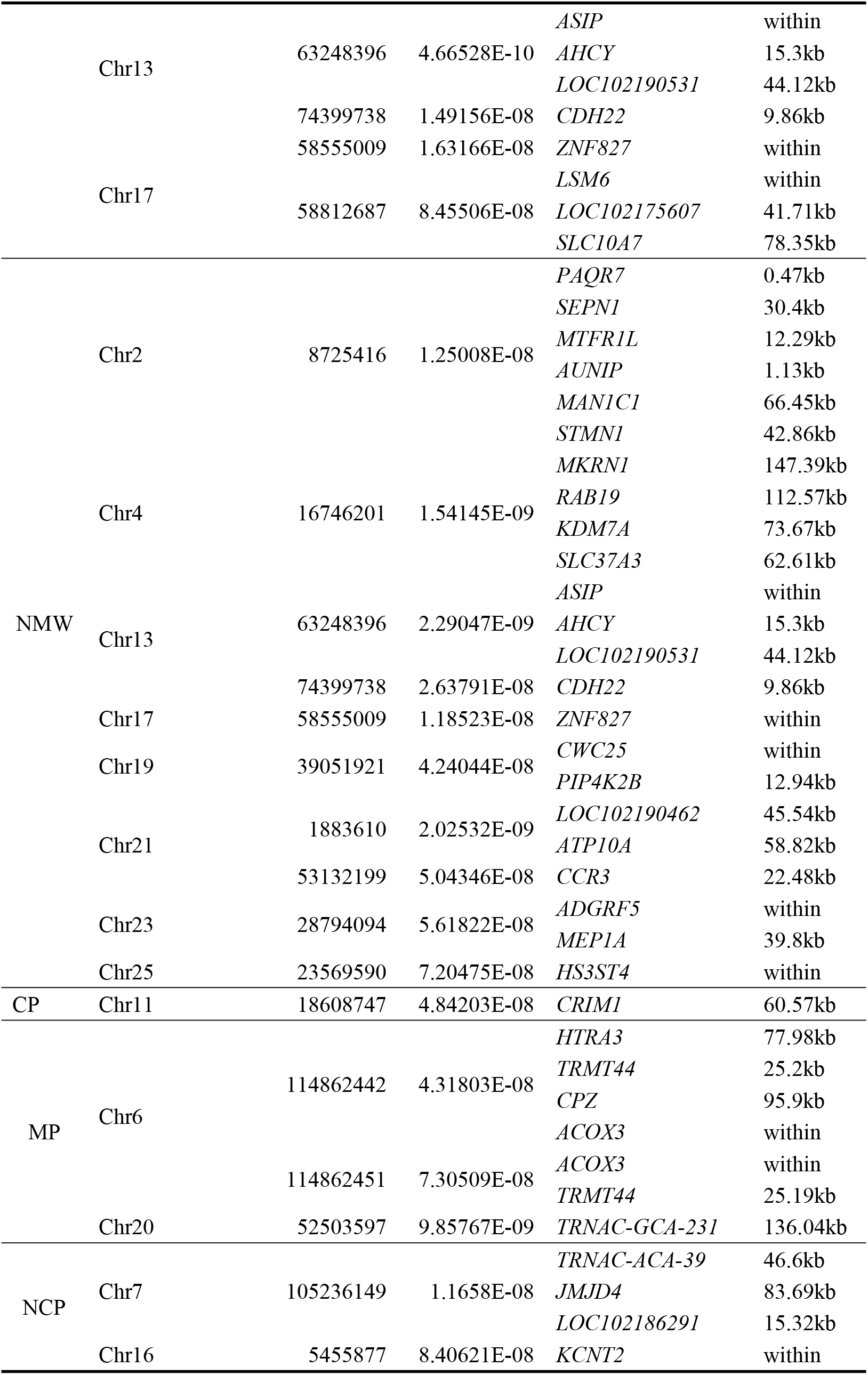
The partial GWAS results of the carcass traits

